# Can a red wood-ant nest be a trap for fault-related CH_4_ micro-seepage? A case study from continuous short-term *in-situ* sampling

**DOI:** 10.1101/154245

**Authors:** G.M. Berberich, A.M. Ellison, M.B. Berberich, A. Grumpe, A. Becker, C. Wohler

**Author notes:** These authors contributed equally to this work.

## Abstract

Methane (CH_4_) is common on Earth, forms the major commercial natural gas reservoirs, and is a key component of the global carbon cycle, but its natural sources are not well-characterized. We present a geochemical dataset acquired from a red wood-ant (RWA; *Formica polyctena*) nest in the Neuwied Basin, a part of the East Eifel Volcanic Field (EEVF), focusing on methane (CH_4_), stable carbon isotope of methane (δ^13^C-CH_4_), RWA activity patterns, earthquakes, and earth tides. Nest gas and ambient air were continuously sampled *in-situ* and analyzed to detect microbial, thermogenic, and abiotic fault-related micro-seepage. Methane degassing was not synchronized with earth tides. Elevated CH4 concentrations in nest gas appear to result from a combination of microbial activity and fault-related emissions moving via through fault networks through the RWA nest. Two δ^13^C-CH_4_ signatures were identified in nest gas: −69‰ and −37‰. The −69‰ signature of δ^13^C-CH_4_ within the RWA nest is attributed to microbial decomposition of organic matter. This finding supports previous findings that RWA nests are hot-spots of microbial CH_4_. Additionally, the −37% δ^13^C-CH_4_ signature is the first evidence that RWA nests also serve as traps for fault-related emissions of CH_4_. The −37‰ δ^13^C-CH_4_ signature can be attributed either to thermogenic/fault-related or to abiotic/fault-related CH_4_ formation originating from e.g. low-temperature gas-water-rock reactions in a continental setting at shallow depths (micro-seepage). Sources of these micro-seeps could be Devonian schists (“Sphaerosiderith Schiefer”) with iron concretions (“Eisengallen”), sandstones, or the iron-bearing “Klerf Schichten”. We cannot exclude overlapping micro-seepage of magmatic CH_4_ from the Eifel plume. Given the abundance of RWA nests on the landscape, their role as sources of microbial CH_4_ and traps for abiotically-derived CH_4_ should be included in estimation of methane emissions that are contributing to climatic change.

## Introduction

Methane (CH_4_) is common on Earth, forms the major commercial natural gas reservoirs, and is a key component of the global carbon cycle [1–2]. This second-most important greenhouse gas currently has an average atmospheric concentration of 1.82 ppm, and continues to increase [3]. Today, most natural occurrences of CH_4_ are associated with terrestrial and aquatic processes. In the shallow subsurface, CH_4_ is produced on geological time scales mainly by thermal conversion of organic matter resulting from heat and pressure deep in the Earth’s crust or by microbial activity. This biotic CH_4_ includes the formation of thermogenic CH_4_ and microbial aceticlastic methanogenesis [4–5]. In contrast, abiotic CH_4_ is produced in much smaller amounts on a global scale and is formed by either high-temperature magmatic processes (Sabatier-type reactions) in volcanic and geothermal areas, or via low-temperature (<100 °C) Fischer-Tropsch-Type (FTT) gas-water-rock reactions in continental settings, even at shallow depths. It is found in specific geologic environments, including volcanic and geothermal systems; fluid inclusions in igneous intrusions; crystalline rocks in Precambrian Shields; and submarine, serpentinite-hosted hydrothermal fields or land-based serpentinization fluids [2,4].

In most geologic environments, biotic and abiotic gases occur simultaneously. Both thermogenic and abiotic CH_4_ reach the atmosphere through marine and terrestrial geologic gas (micro-)seeps, and during the exploitation and distribution of fossil fuels. To identify whether locally elevated CH_4_ concentrations in the atmosphere are due transportation via fault networks, a determination of possible methane source(s) is required. At the land surface, CH_4_ is produced by methanogenic Archaea in anaerobic soil environments or through oxidation by methanotrophic bacteria in aerobic topsoils [6]. Isotopic measurements of δ^13^C-CH_4_ can distinguish abiotic from biotic CH_4_ [7–8].

Increase in compressive stress, changes in the volume of the pore fluid or rock matrix, and fluid movement or buoyancy are important mechanisms driving fluid flow and keeping fractures open [9–10]. Faults and fracture networks act as preferential pathways of lateral and vertical degassing, creating linear fault-linked anomalies, irregularly-shaped diffuse or “halo” anomalies and irregularly-spaced plumes or “spot anomalies” [e.g. 11–12]. [10] showed that faults had δ^13^C-CH_4_ = −37%_0_ and a significantly higher CH_4_ flux (11.5±6.3 t CH_4_ km^−1^ yr^−1^) than control zones. In Europe, micro-seeps occur both onshore and offshore, with estimated CH_4_ flux in Europe of 0.8 Tg yr^−1^ and total seepage of 3 Tg yr^−1^ [12, 5].

Recent research has revealed close relationships between the spatial distribution of red wood-ant nests (*Formica rufa*-group; henceforth RWA) and tectonic fault zones [13–16]. Exploratory testing of fault-zone gases revealed that helium (He) and radon (Rn) in RWA nests exceeded atmospheric and background concentrations [13–14]. RWA mounds also have been found to be “hot spots” for CO_2_ emissions in European forests [17–20]. [21] showed that ant mounds (*Lasius flavus, Lasius niger* and *Formica Candida*) contributed measurable amounts to soil gas emissions from wetlands (CO_2_: 7.02% and N_2_O: 3.35%), but act as sinks with regard to the total soil CH_4_ budget (-4.28%). In contrast to that, higher net CH_4_ emission (3.5 μg m^2^h^−1^) were found in fire ant mounds situated in natural pasture soils [22]. However, continuous *in-situ* sampling of natural release of CH_4_ from RWA nests has not been done. [6] estimated CH_4_ flux from two nest material samples collected from the top and the rim of each of five nests on two different days (30 July and 14 October) in 2014. Finally, natural release of CH_4_ via fault zones [10] has been rarely considered, although there are a range of processes that could contribute to it, including micro-seepage via buoyant flux of CH_4_, faults increasing the flow rate of microbubbles, and gas vents responding to earth tides and earthquakes [23–24].

We used a combination of geochemical, geophysical, and biological techniques; state-of-the-art image analysis; and statistical methods to identify associations between RWA activity, continuous *in-situ* CH_4_ degassing, earth tides, and tectonic processes. We aimed to test from a geochemical/geophysical point of view three different hypothesis: a) whether a RWA nest can indicate actively *in-situ* degassing faults trapping migrating CH_4_ from the deep underground; b) whether RWA activity changes during the CH_4_ (micro)-seepage process; and c) whether CH_4_ (micro)-seepage process is affected by external agents (earth tides, earthquake events, or meteorological conditions). Specifically, we tested the null hypotheses that, in the field, *in-situ* concentrations of both CH_4_ and δ^13^C-CH_4_ and RWA activity are independent. We found a RWA nest appears to trap fault related micro-seepage of CH_4_, and that degassing pattern are independent from earth tides and meteorological conditions.

## Methods

We explored associations between RWA activity, *in-situ* methane concentrations in an ant nest and ambient air, tectonic events, and weather processes and earth tides at the Goloring site near Koblenz, Germany during a continuous, *in-situ* 8-d sampling campaign that ran from 4-11 August 2016. Time and duration of the CH_4_ sampling was determined by the availability of the CRDS analyser owned by the Institute for Geosciences (University of Heidelberg). Our approach contrasts with prior work where different nests were statically sampled and CH_4_ flux was estimated from 10 nest-material samples [e.g. 6].

### Study area

The Goloring site is located west of the Rhine River, southeast of the Laacher See volcano, and close to the Ochtendung Fault Zone in the seismically active Neuwied Basin, which is part of the Quaternary East Eifel Volcanic field (EEVF; western Germany; Fig 1A). The EEVF includes ≈100 Quaternary volcanic eruption centers; the Laacher See volcano experienced a phreato-plinian eruption ≈12,900 years ago [25]. The Paleozoic basement consists of alternating strata of Devonian, iron-bearing, quartzitic sandstones with a carbonate matrix and argillaceous shale reaching to 5-km depths. Several thin black coal seams (Upper Siegen) are embedded within these alternating strata [26]. Ecocene/Oligocene lignite seams are found at ≈75-160 m and are covered by Paleogene volcanites and Neogene clastic sediments. The study area has been affected by complex major tectonic and magmatic processes, including plume-related thermal expansion of the mantle-lithosphere [27–29], crustal thinning and associated volcanism [30], active rifting processes [31], and possibly crustal-scale folding or the reactivation of Variscan thrust faults under the present-day NW-SE-directed compressional stress field [31–32]. Those processes can be attributed to the existence of old zones of weakness that are reactivated under the current stress field [33–34,29]. Earthquakes (Fig 1A) are concentrated in areas that are related to the seismically active Ochtendunger Fault Zone [33]. These earthquakes are related to stress-field-controlled block movements, have a weak-to-moderate seismicity, and occur mostly in a shallow crustal depth (≤15 km) with local magnitudes (Richter scale) rarely exceeding 4.0. No fault zones have been reported from our Goloring study site, and focal depth of earthquakes near the site never exceeded 28 km during our sampling campaign [35].

**Fig 1.**
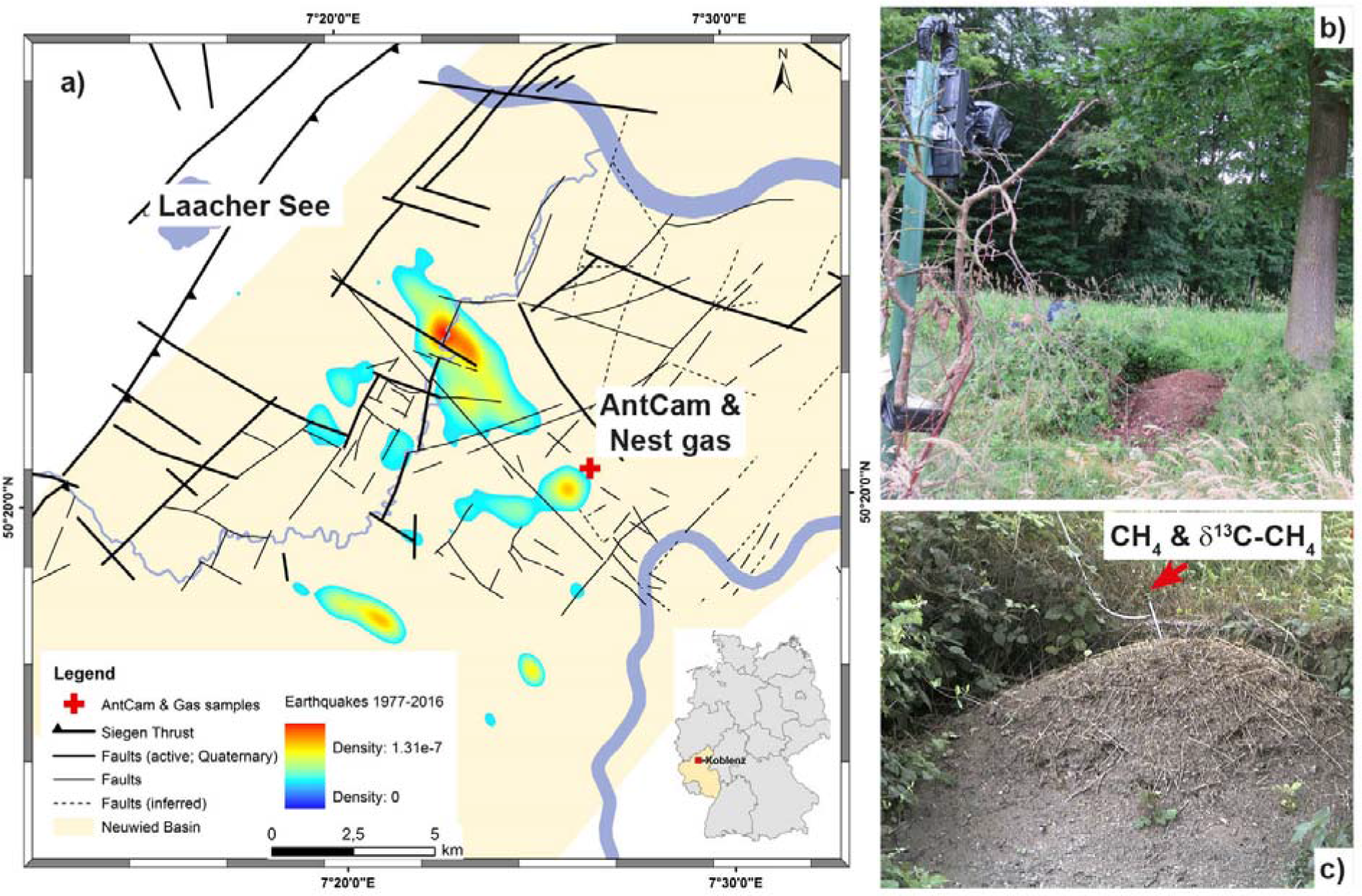
Location of the Goloring study area within the Neuwied Basin. The map (a) shows the Goloring study site (red cross) ≈15 km SE of the Laacher See volcano within the Neuwied Basin (light yellow area), tectonic structures (black lines) and probability density of the earthquake events from 1977-2016 which are related to the Ochtendunger Fault Zone (rainbow contours). The inset shows the location of study site within Germany. Photographs show (b) the AntCam for continuous monitoring of ant activity and (c) the nest gas probe (all photographs: G. Berberich).

### Monitoring red wood ant activity

Within the research project “GeoBio-Interactions” (March – September, 2016), we monitored RWA activity using an “AntCam”: a high-resolution camera system (Mobotix MX-M12D-Sec-DNight-D135N135; 1,280 × 960 pixels) installed ≈5 m from a RWA nest (Fig 1B). During the 192-hr CH_4_ sampling campaign, which ran from 4-11 August, 2016, ant activities were recorded and time-stamped continuously (12 Hz). The network-compatible AntCam was connected to a network-attached storage (NAS) system for data storage via a power-over-Ethernet (POE) supply. A computer connected to the NAS evaluated the RWA activities on-site and in real time using C++ code to accelerate image evaluation. Image analysis extended the system of [36] and was based on the difference image technique (Fig 2). To reduce negative influences caused by, e.g., moving blades of grass, we used a mask to restrict analysis to only the visible top of the mound. To compensate for slight movements of the camera, e.g., due to wind, an image registration of the current image relative to the previous image was done based on mutual information before the determination of the absolute difference image [37]. Results of RWA activity were written to a file. Every hour, this file was sent via email (mobile data transfer, LTE router) to a mail server. Since two different sensors were used for the day and night, respectively, we computed different polynomials to map the sum of absolute differences onto manually designed activity categories in a follow-up procedure. The coefficients of the polynomials were obtained from a minimization of the sum of squared differences between the polynomial model and the manually assigned category for two selected weeks. A first-order polynomial was adapted to the daytime data and a third-order polynomial was adapted to the nighttime data. To avoid numerical difficulties, we first centered and scaled the data by subtracting the mean of the data during the target time and dividing by the standard deviation. Both values were computed for day-and nighttime, respectively.

**Fig 2.**
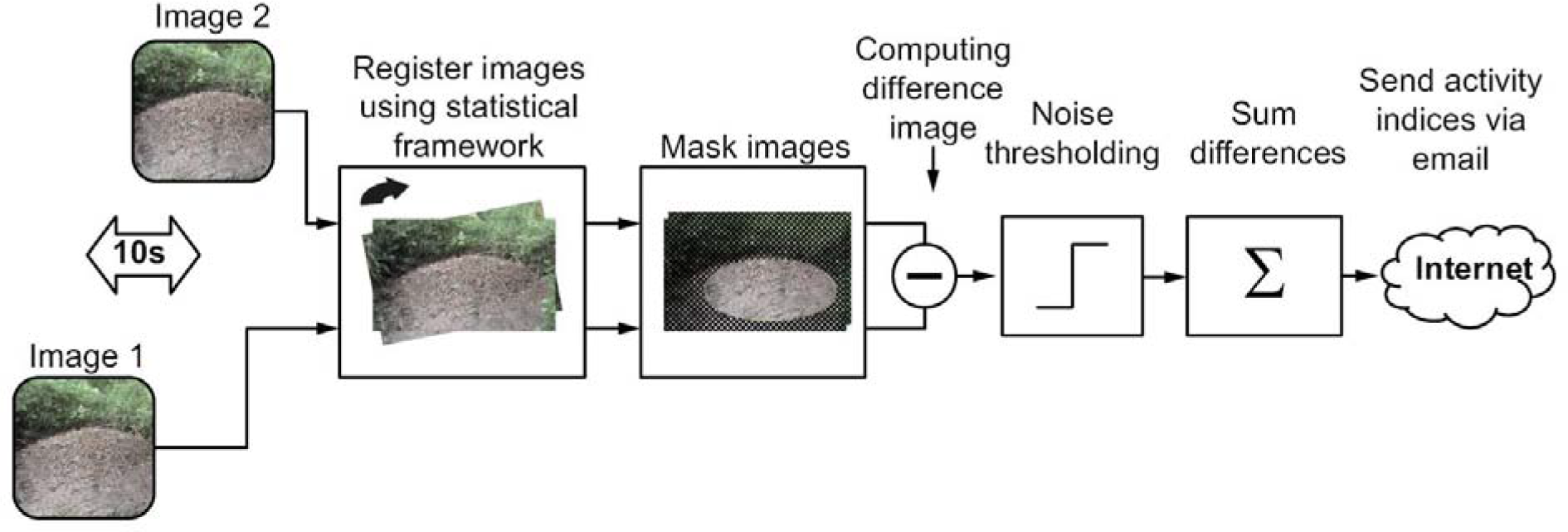
Workflow for acquisition and estimation of RWA activity. The “AntCam”, a network-compatible high-resolution camera system, was connected to a network-attached storage (NAS) system for data storage via a power-over-Ethernet (POE) supply. A computer connected to the NAS evaluated the RWA activities on-site and in real time using C++ code to accelerate image evaluation. Image analysis extended the system of [36] and was based on the difference image technique. Results of RWA activity were written to a file. Every hour, this file was sent via email (mobile data transfer, LTE router) to a mail server.

### Gas sampling and geochemical analyses

Field measurements of CH_4_ were taken from 4–11 August 2016. A stainless-steel probe (inner diameter 0,6 cm; Fig 1C) was inserted into the *F. polyctena* nest to a depth of 80 cm and remained there, unmoved, during the entire 192-hour sampling campaign. The probe was used for continuous CH_4_/δ^13^C-CH_4_ measurements. The probe was equipped with a flexible tip attached to a pushable rod and a sealable outlet for docking sampling equipment. The closed probe was inserted into the nest. After opening by pushing the rod, the probe was evacuated twice, using a 20-ml syringe. After this, the outlet was closed to prevent atmospheric influence. The outlet was only opened after docking the sampling unit to it.

Concentrations of CH_4_ and δ^13^C-CH_4_ in nest gas (NG) and ambient air (AA) were monitored using a portable CRDS analyser (G2201-i; Picarro, USA) that measured ^12^CH_4_, ^13^CH_4_ and H_2_O quasi-simultaneously at 1 Hz, and provided δ^13^C values relative to the Vienna Pee Dee Belemnite standard. The G2201-i uses built-in pressure and temperature control systems, and automatic water-vapor correction to ensure high stability of the portable analyzer. Effects of water vapor on the measurement were corrected automatically by the Picarro^®^ software. The manufacturer guarantees concentration precision for the analysis of CH_4_ in the “high precision mode” of 5 ppbv ± 0.05 % (^12^C) and 1 ppbv ± 0.05% (^13^C) within a concentration range of 1.8-1000 ppm. The guaranteed precision of δ^13^C-CH_4_ is <0.8‰.

The CRDS analyzer was deployed in a dry, wind-sheltered location near the RWA nest. Nest gases were pumped from the aforementioned probe into the CRDS analyzer for analysis of CH_4_ and δ^13^C-CH_4_ values. Ambient air was measured 2 m away from the nest for 15 min every four hours during the operation using a 3-way-valve, avoiding disturbance of the nest or the position of the steel probe. All gases passed through a chemical trap filled Ascarite^®^ (sodium hydroxide coated silica; http://www.merckgroup.com) before entering the system to remove carbon dioxide (CO_2_) because the high concentrations of CO_2_ in the nest samples could interfere with the measurements of CH_4_ and δ^13^C-CH_4_. Gas samples were dried by a Nafion^®^ drying tube (Nafion MD110, PermaPure LLC, USA) before measurements to ensure higher accuracy and subsequently analyzed for CH_4_ concentration and δ^13^C-CH_4_. To assure quality of the CH_4_ and δ^13^C-CH_4_ values, reference gas measurements were taken every 8 h during the operation. Fluctuations in atmospheric CH_4_ and δ^13^C-CH_4_ values were validated against a single, 4-h measurement of ambient air. Carbon isotope ratios are expressed using standard delta (6) notation as described by deviations from a standard: δ_sample‰_ = ((R_sample_/R_standard_-1)) × 1000, where R is the ^13^C/^12^C ratio in the sample or standard. A total of 459 704 samples for both CH_4_ and δ^13^C-CH_4_ in nest gas and 27 samples in ambient air were collected and analyzed.

### Meteorological Parameters

A radio meteorological station (WH1080) placed 2 m above the ground at the Goloring site continuously logged meteorological conditions (temperature [°C], humidity [%], air pressure [hPa], wind speed [m/s], rainfall [mm], and dew point [°C]) at 5-min intervals. The recorded data were downloaded every two days, checked for completeness, and stored in a data base.

### Earth tides

Cyclic changes in the earth’s environment are caused by the gravitational pull of both the Sun and the Moon on the earth. These result in two slight lunar and two solar tidal bulges (“earth tides”). The two bulges occur at the surface of the earth that approximately faces the Moon and at the opposite side while the Earth rotates around its axis. Earth tides were calculated using the tool developed by [38].

### Earthquake events

Data on earthquake events during the sampling campaign were obtained from the seismological databases provided by the Erdbebenstation Bensberg [35, http://www.seismo.uni-koeln.de/events/index.htm] and by the Landesamt für Geologie und Bergbau, Rheinland-Pfalz [39, http://www.lgb-rlp.de/fachthemen-des-amtes/landeserdbebendienst-rheinland-pfalz/]. The probability density of the earthquake events was estimated using the kernel density estimator of [40] using Gaussian kernels.

### Data analysis

All analyses were done using R version 3.3.2 (R Core Team 2016) or MATLAB R2017a.

We examined associations between the six measured meteorological variables and RWA activity and CH_4_ concentrations. As many of these variables were correlated with one another, we used principal components analysis (R function prcomp) on centred and scaled data to create composite “weather” variables (i.e., principal axes) that were used in subsequent analyses.

We used the “median+2MAD” method [41] to separate true peaks in CH_4_ concentrations from background or naturally-elevated concentrations: any observation greater than the overall median+2MAD (2.31 ppm CH_4_ in nest gas and 2.11 ppm CH_4_ in ambient air) was considered to be a peak concentration. Background and elevated CH_4_ concentrations were separated based on the 90% quantile of the CH_4_ concentration [42]. For interpreting the significance of the correlation coefficient, we followed [43]. For δ^13^C-CH_4_, we considered concentrations < −35‰ or > 0‰ to be peak concentrations. Only peaks occurring in both data sets at the same time were considered to be true peaks. The Keeling plot method [44] was applied to determine the carbon-isotope composition of the found peaks to obtain insights into the processes that govern the distinction between isotopes in the ecosystem.

## Availability of data

Data are available from the Harvard Forest Data Archive (http://harvardforest.fas.harvard.edu/data-archive), dataset HF-XXX.

## Results

### Meteorological conditions

During the one-week field campaign in August 2016, air temperatures ranged from 5.7-29.1 °C (mean = 16.2 °C), with only 2.1 mm rainfall overnight between 9 and 10 August. Variation in atmospheric pressure (mean 988 ± 2.24 hPa) and wind speed (1.67 ± 1.72 km/h) were small. The first three axes derived by the principal components analysis accounted for nearly 80% of the variance in the data (Table 1). The first axis represents temperature and humidity, the second axis represents atmospheric pressure (with additional contributions of humidity and windspeed), and the third axis represents rainfall and windspeed (with a minor contribution of temperature).

**Table 1.** **Results of the principal components analysis of the measured weather variables. Values in the first six rows are the loadings of each variable on each of the first three principal axis; only loadings > |0.3| are shown. The last row of the table gives the cumulative proportion of the variance explained by each of the first three principal axes**.

| Variable | PC-1 | PC-2 | PC-3 |
| --- | --- | --- | --- |
| Temperature (°C) | −0.69 |  | 0.3 |
| Atmospheric Pressure (hPa) |  | 0.62 |  |
| Dew-point (°C) | −0.43 | 0.49 |  |
| Relative humidity (%) | 0.48 | 0.45 |  |
| Rainfall (mm) |  |  | 0.77 |
| Windspeed (km/h) |  | −0.41 | 0.50 |
| **Cumulative variance explained** | **0.34** | **0.59** | **0.77** |

Median RWA activity and the three principal axes of weather were modestly associated, and accounted for only 8% of the variance in ant activity (Table 2). The ant activity increased slightly at lower temperatures (PC-1) and slightly decreased when rainfall (PC-3) was present. PC-2 was not associated significantly with RWA activity.

**Table 2.** **Summary ANOVA table of the linear model examining the effects of weather conditions on median RWA activity. The estimate is the slope describing the relationship between each principal component and median RWA activity. The remaining columns are the degrees of freedom, mean square, and F-statistic for each term in the model. (\*\*\**P* < 0.001; ^NS^*P* > 0.5). Overall model *r*^2^ = 0.08; F_3,1536_ = 47.57, *P* < 0.001**.

|  | Estimate | Df | MS | F |
| --- | --- | --- | --- | --- |
| PC-1 | 0.19 | 1 | 110.8 | 120.6*** |
| PC-2 | -0.01 | 1 | 0.3 | 0.3<sup>NS</sup> |
| PC-3 | -0.11 | 1 | 20.0 | 21.8*** |
| Residual |  | 1563 | 0.9 |  |

Weather conditions explained 10% of the variation in CH_4_ (ppm) (Table 3), but explained 22% of the variation in δ^13^C-CH_4_ (‰), which decreased with all measured weather variables (Table 4).

**Table 3.** **Summary ANOVA table of the linear model examining the effects of weather conditions on CH_4_ concentration (ppm). The estimate is the slope describing the relationship between each principal component and CH_4_ concentration. The remaining columns are the degrees of freedom, mean square, and F-statistic for each term in the model. (\*\*\**P* < 0.001; \**P* < 0.5). Overall model *r*^2^ = 0.19; F_3,1536_ = 121.5, *P* < 0.001**.

|  | Estimate | Df | MS | F |
| --- | --- | --- | --- | --- |
| PC-1 | -0.3 | 1 | 263.1 | 323.8*** |
| PC-2 | 0.1 | 1 | 29.8 | 36.6*** |
| PC-3 | 0.04 | 1 | 3.4 | 4.1* |
| Residual |  | 1563 | 0.8 |  |

**Table 4.**
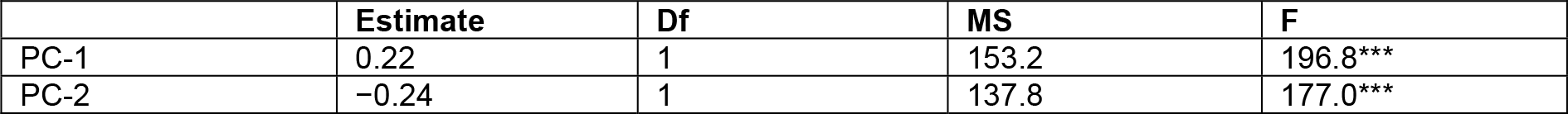
**Summary ANOVA table of the linear model examining the effects of weather conditions on δ^13^C-CH_4_ (‰). The estimate is the slope describing the relationship between each principal component and CH_4_ concentration. The remaining columns are the degrees of freedom, mean square, and F-statistic for each term in the model. (\*\*\**P* < 0.001). Overall model *r*^2^ = 0.22; F_3,1536_ = 149.5, *P* < 0.001**.

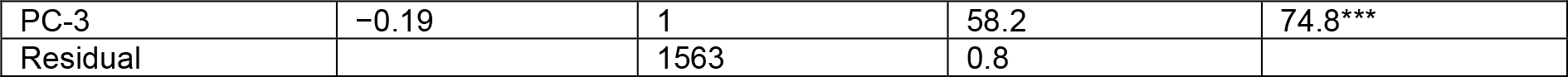

### RWA activity

Ants were most active during the late afternoon and early evening hours (Fig 3A; 4A). The video streams showed that the ants went on foraging, building and maintaining the nest as they had done since the start (on March, 18^th^) of our longer 7-month field campaign. Decomposition of the time-series into its additive components (Fig 3B-D) illustrated that during the one-week gas-sampling campaign, there was a trend towards increasing activity over the first four days, followed by a sharp decline towards the end of the week (Fig 3B). There were two noticeable peaks of activity, at mid-day and early afternoon, followed by sharp spikes in activity near 16:30 hours (Fig 3C).

**Fig 3.**
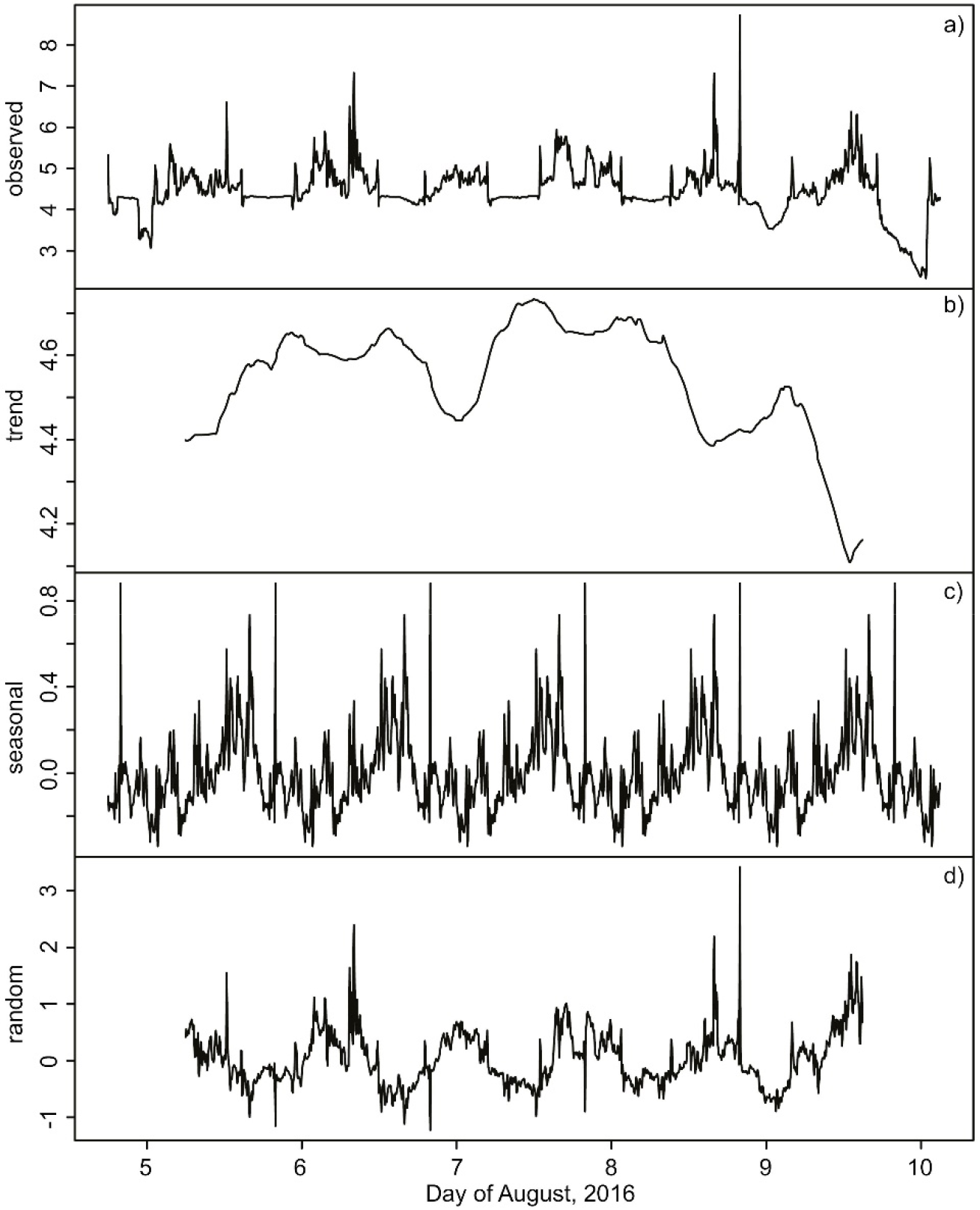
Additive time-series decomposition of median RWA activity. An extreme spike in ant activity (observed = 12 units on 04-August at 19:14 UTC and 25 units on 04-August at 19:19 UTC) are not shown to enhance clarity of the “observed” time-series. The observed data (**a**) can be partitioned (additively decomposed) into its temporal trend estimated by a polynomial smoothing function (**b**), a daily (“seasonal”) cycle (**c**); and residual (random) variation (**d**).

Additional external agents that may have influenced RWA activity were also visually assessed: No nuptial flight happened during this week. Ventilation phases of the nest took place in the early morning (6:40 - 7:30 UTC) on 5 August for 50 minutes and on 7 August for 20 minutes (6:40 - 7:00 UTC) after sunrise with varying ant activities (Fig 4A). On two days (07.08. and 09.08.), at 04:30 and 05:50 (UTC), respectively, golden hammer birds (*Emberiza citronella*) were “anting” for ≈5 min to kill parasites on their feathers with formic acid; a mouse was observed on the nest at 22:00 (UTC) for 10 minutes on 04.08.16. These biotic effects did not appear to influence any RWA activity.

**Fig 4.**
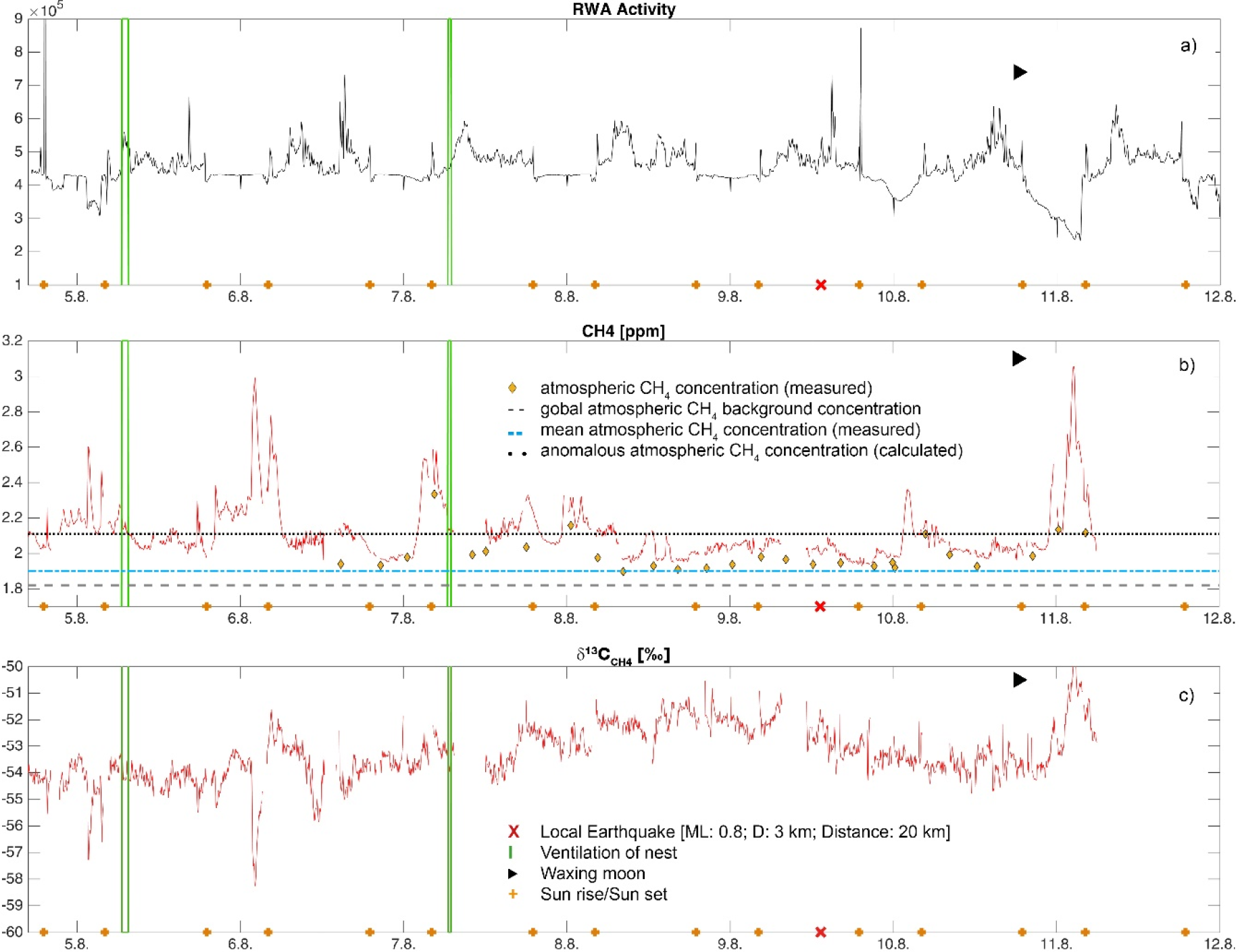
Time-series plots of median RWA activities (a), CH_4_ (b), and δ^13^C-CH_4_ (c) in nest gas. Green lines indicate ventilation phases of the nest, orange crosses sunrise/sunset, and a red cross a single local earthquake. Reference lines indicate the global atmospheric CH_4_ background concentration (Saunois et al. 2016; black dashed line), the local mean CH_4_ atmospheric concentration (blue dotted line), and the calculated anomalous atmospheric CH_4_ concentration (black dotted line) during the sampling week in August 2016.

### CH_4_ and δ^13^C-CH_4_ in nest gas

A total of 459,704 data points were collected during the 192-hr sampling for each of CH_4_ and δ^13^C-CH_4_. Concentrations of CH_4_ in the nest exceeded the global atmospheric background concentration (1.82 ppm; Saunois et al. 2016) and ranged from 1.93 to 3.07 ppm (Fig 4B, Table 5). Atmospheric CH_4_ concentrations were slightly variable (1.90 - 2.33 ppm). The calculated anomalous threshold concentration after [41] for atmospheric CH_4_ was 2.11 ppm CH_4_ (Fig 4B). In ambient air, only four measurements out of 27 exceeded this threshold. In nest gas, the anomalous threshold was 2.31 ppm CH_4_. To compare our findings to fault-related emissions [10], the 90^th^ percentile of CH_4_ was estimated. In nest gas, 10% of measured CH_4_ was larger than the 90^th^ percentile (Table 5). Nest gas concentrations of CH_4_ appear to result from fault-related emissions moving via fault networks through the RWA nest. A comparison with fugitive emissions of CH_4_ (ppm) from basin bounding faults in the UK [10; Table 5] showed that mean nest gas emissions are of the same order, although we had 20× more observations.

**Table 5.**
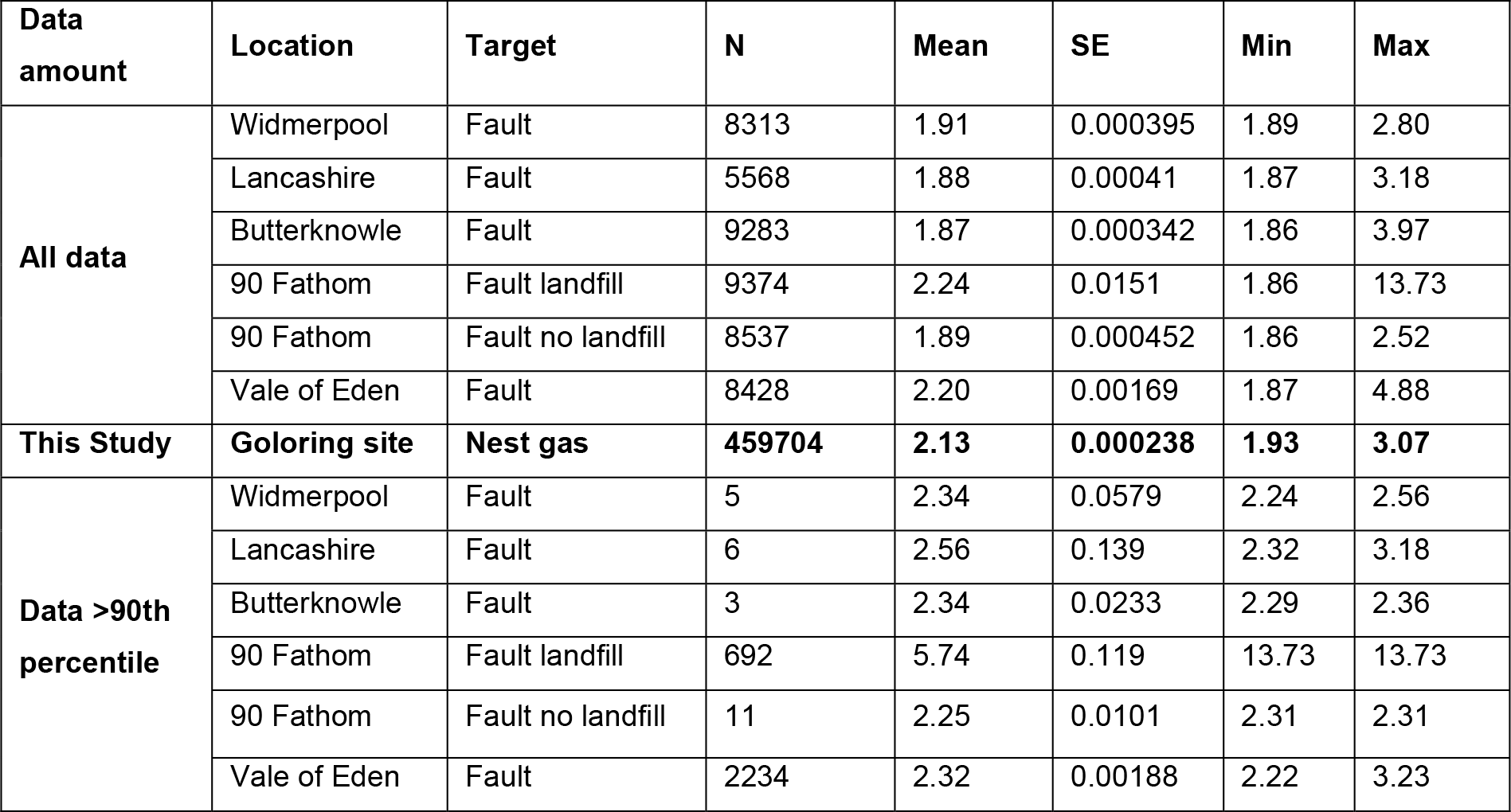
**Descriptive statistics for nest gas CH_4_ (ppm) at the Goloring site compared to fugitive emissions of CH_4_ (ppm) from basin bounding faults in the UK [10]. SE = 1 standard error of the mean**.

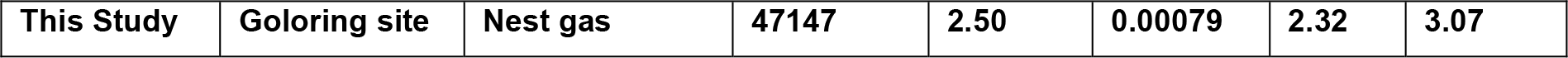

δ^13^C-CH_4_ in the nest ranged from −58.48 to −49.54‰ (Fig 4C). Eight significant peaks (red and blue marks in Fig 5A, B) in nest gas were found for CH_4_ and δ^13^C-CH_4_ (Fig 5a, b). These peaks occurred between 17:39 (UTC) and 06:54 (UTC) the following day, but were otherwise not temporally predictable. Results of the Keeling plots [44] revealed two signatures for δ^13^C-CH_4_ at −37‰ (blue markers and dots in Fig 5A, B and C) and −69‰ (red markers and dots in in Fig 5A, B and C) in nest gas (Fig 5C).

**Fig 5.**
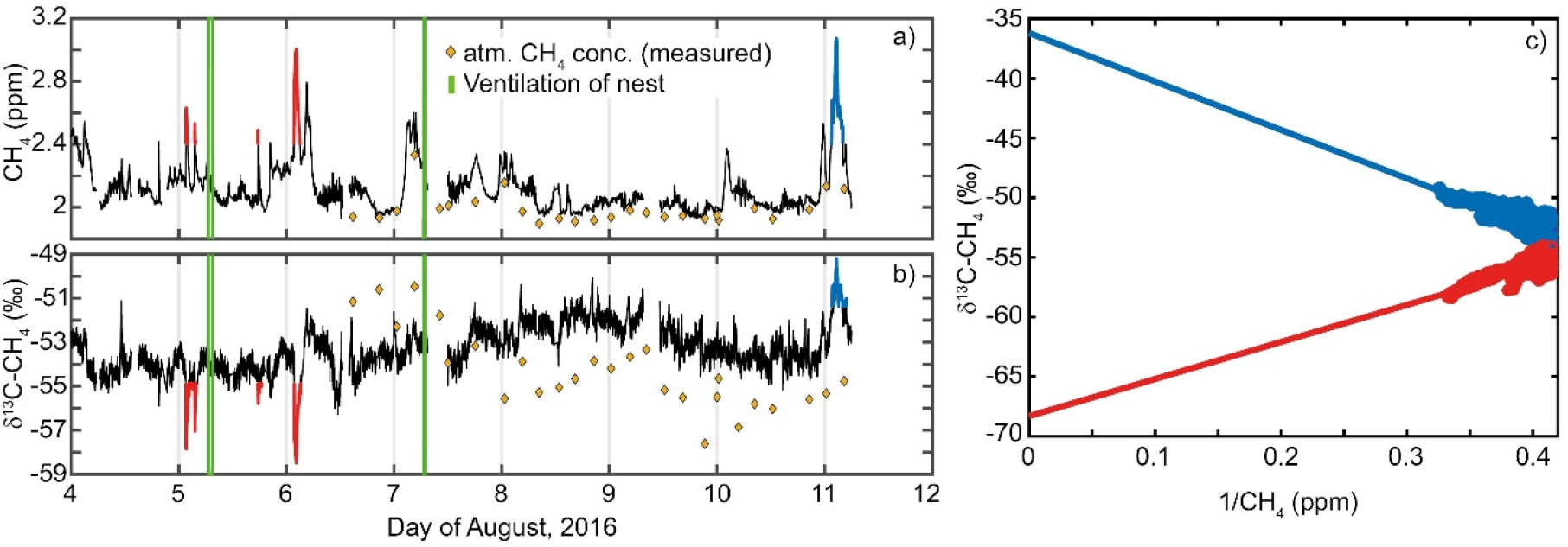
CH_4_ (a), δ^13^C-CH_4_ (b) peak concentrations and Keeling plot of δ^13^C-CH_4_ (c) from nest gas. Note the peaks indicate two signatures for δ^13^C-CH_4_ in nest gas at −37‰ and −69‰ (c). For better identification of the signatures in the Keeling plot, peak concentrations in CH_4_ and δ^13^C-CH_4_ were colored in a) and b). Red signatures in the Keeling plot refer to the marked red peak concentrations in a) and b), whereas blue signatures in the Keeling plot to the marked blue peak concentrations in a) and b).

Joint visualization of the time series of ant activity, methane concentrations, and weather (Fig 6A) reveal that all the time series exhibited a periodicity of approximately 24 hours. Cross-correlations showed positive and negative peaks at daily intervals (Fig 6B). The absolute value of the crosscorrelation coefficient ≤ 0.3, and the strongest cross-correlation occurred at a lag of ≈-30 minutes, less than the original filter width of the ant activity time series.

**Fig 6.**
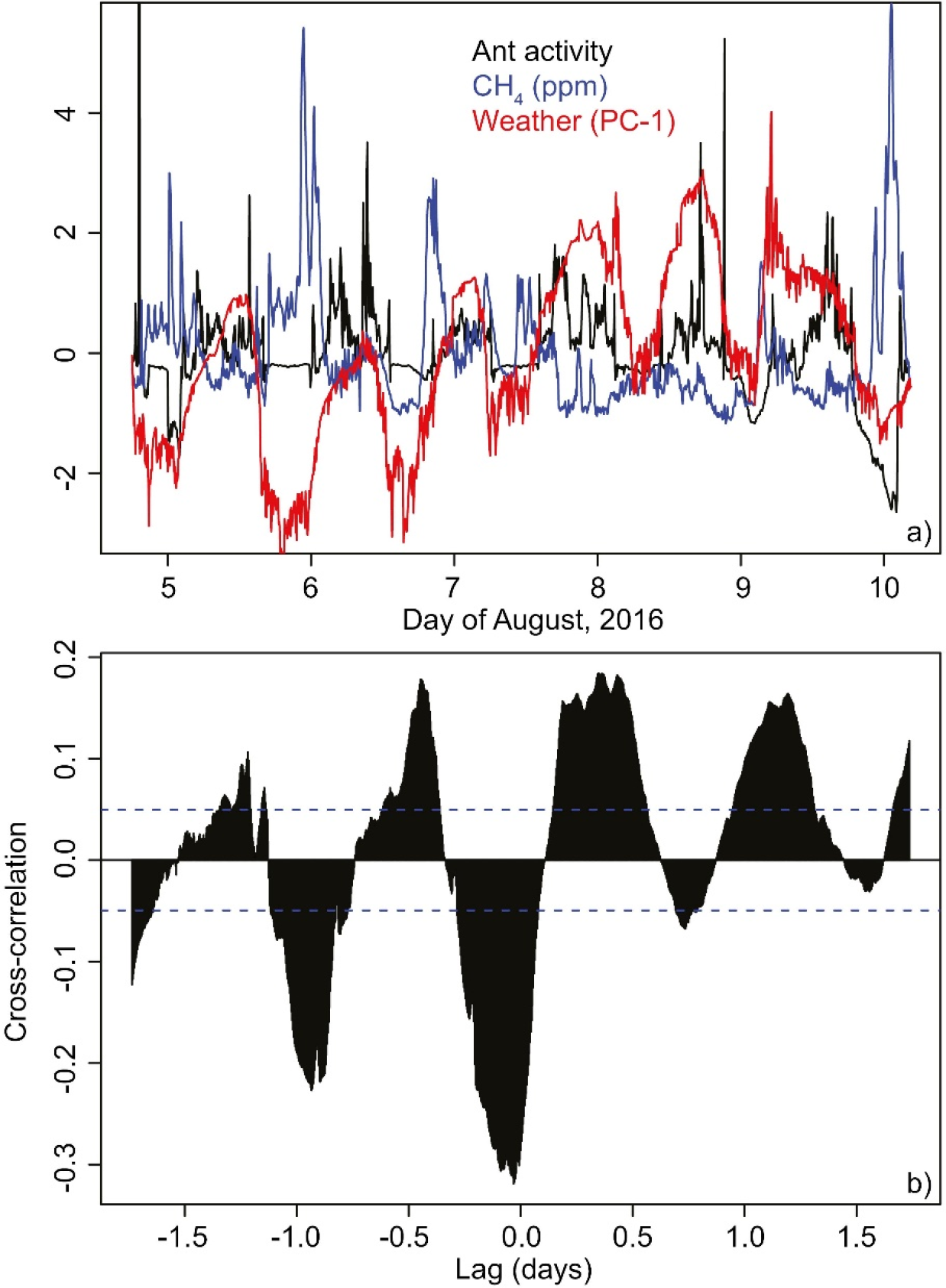
Relationships between ant activity, CH_4_ and weather conditions. Time-series plot (a) of median ant activity (black), methane concentration (blue), and weather conditions (PC-1, red). Cross-correlation (b) between median ant activity and methane degassing. All values are centered and scaled (i.e., are reported in SD units).

### Earthquakes

Only one earthquake [35,39] occurred nearby (local magnitude: 0.8; depth: 3 km; distance: 20 km; Fig 4). This micro-earthquake neither influenced degassing nor RWA activity.

## Discussion

Our results provide for the first time a continuous *in-situ* record of 192-hr sampling of both CH_4_ and δ^13^C-CH_4_ in a RWA nest. Although our results of CH_4_ and δ^13^C-CH_4_ in nest gas may not be representative of these values for the entire year, the measurement data provide a continuous set of observations of multiple variables matched in time, in contrast to other data reported in literature for which different nests were sampled at different times (two days) and CH_4_ flux was estimated in lab incubations from 10 nest material samples [e.g. 6].

### CH_4_ and δ^13^C-CH_4_ in nest gas

Results from our short but continuous *in-situ* sampling confirmed our 1^st^ hypothesis that elevated CH_4_ concentrations in nest gas appear to result from fault-related emission moving via fault networks through the RWA nest. In contrast to [21] our results show that also a red wood ant nest acts as a CH_4_ source. [22] attribute nest gas CH_4_ to high NH4-N concentrations in ant mounds. A comparison of our results with data on fugitive emissions of CH_4_ (ppm) from basin bounding faults in the UK (Boothroyd et al. 2017; Table 5) showed that mean nest gas emissions are of the same order. Elevated CH_4_ concentrations in nest gas appear to result from a combination of microbial activity and fault-related emissions moving via through fault networks through the RWA nest.

Comparison of δ^13^C-CH_4_ nest gas signatures with published data suggests that it can be attributed to two different sources (Fig 7). The δ^13^C-CH_4_ signature of −69‰ in nest gas indicates a microbial source, such as decomposing organic matter that is high in nutrients [8]. This result supports the findings of [22] that the aboveground parts of ant nests are hot-spots of CH_4_ production.

**Fig 7.**
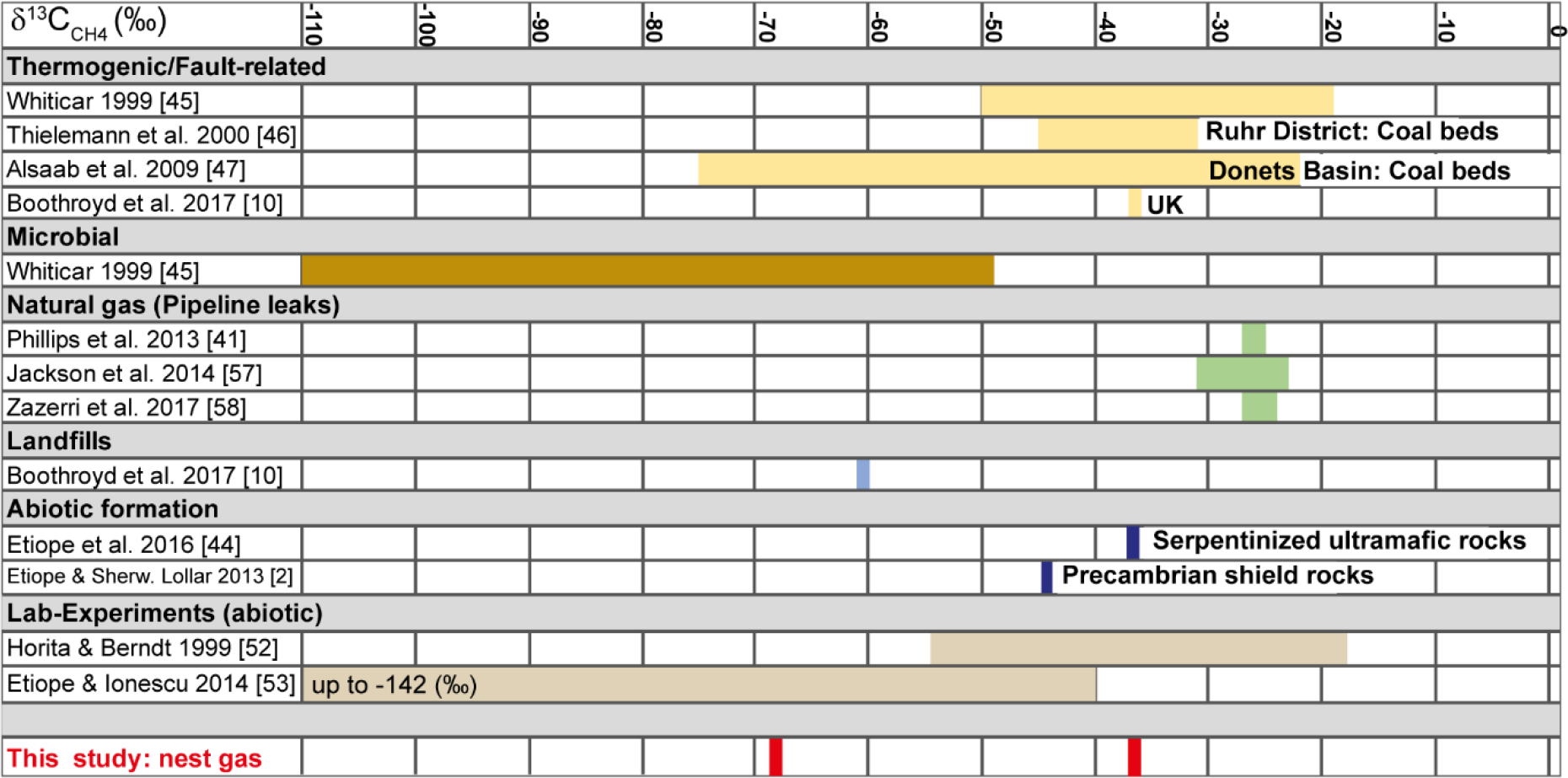
Comparison of δ^13^C-CH_4_ in nest gas signatures to published data. The two nest-gas signatures indicate a microbial source and a thermogenic or abiotic fault-related one.

The second isotope signature, −37‰ δ^13^C-CH_4_, can be attributed either to thermogenic/fault-related [10] or to abiotic/fault-related CH_4_ formation [45]. This result provides the first evidence that RWA nests may serve as traps for fault-related emissions of CH_4_. [10] found a δ^13^C-CH_4_ signature of −37‰ for fugitive emission of CH_4_ via migration along fault zones in the United Kingdom. Our result of −37‰ δ^13^C-CH_4_ is of the same order (Fig 7) and can be attributed to fault-related CH_4_ emission moving through the RWA nest.

Continental loss of volatiles requires tectonically active parts and the formation of fluid-filled conduits through the continental crust. Suitable locations can be found in extensional regimes and their related volcanism [30], such as are present in our study area. Gas permeable faults and fractured rocks are pathways to naturally release significant amounts of "old" CH_4_ of crustal origin. Significant geologic CH_4_ emissions, comprising both biogenic and thermogenic CH_4_, are due to hydrocarbon production in sedimentary basins and, subordinately, to inorganic Fischer-Tropsch type reactions occurring in geothermal systems [24]. A variety of geological, chemical and biological processes have impacts on the deep carbon cycle. There are three possible sources for the fault-related CH_4_ we find in RWA nests.

First, carboniferous coals are sources of thermogenic coalbed methane (CBM) in numerous basins, including the Ruhr and Donets Basins. Their δ^13^C are values between −20‰ and −75‰ [46–48; Fig 9). Both basins have coal thicknesses of ≈100 m [48–49]. In our study area, much older Devonian coal seams with very small thicknesses [26] are reported at depths up to 9000 m. Though the study area is situated in a suitable tectonic compression/extensional regime, any thermogenic CH_4_ would likely be small because of the very low thickness of the seams and might not even lead to measurable coal-bed CH_4_ concentrations in nest gas. On the other hand, lignite and coal formations are often associated with aerobic methylotrophs at depths of over 1 km and are usually considered to be anaerobic [50–52]. In the study area, several small lignite seams (Middle to Upper Eocene) with a thickness of up to 5 m are found in depths of approx. 75 to 160 m. The low thickness and the shallow depth of the lignite may not lead to thermogenic CH_4_ seepage.

Second, δ^13^C-CH_4_ in land-based serpentinized ultramafic rocks can be as light as −37‰, and methane from Precambrian shields may exhibit even lower values (−45‰) [2,4,45]. Laboratory experiments have produced abiotic methane with a wide range of δ^13^C-CH_4_ signatures, including isotopically “light” values once thought to be indicative of biological activity (e.g. −19 to −53.6‰ by [53]; −41 to −142‰ by [54]). Abiotic CH_4_ can be mistaken for biotic CH_4_ of microbial or thermogenic origin because minor amounts of abiotic gas in biotic gas may prevent its recognition based on C and H isotope analysis [55, 45]. Sources of abiotic CH_4_ formation in the study area can be attributed to magmatic CH_4_ formation due to late magmatic (<600°C) re-distribution of C-O-H fluids during magma cooling or gas-water-rock-interactions even at low temperatures and pressures [2]. In the study area, the magmatic source for magmatic CH_4_ formation could be the so called “Eifel plume”, a region of about 100-120 km in diameter between 50-60 km depth and at least 410 km depth beneath the study area. The buoyant Eifel plume is characterized by excess temperature of 100-150 K, has approx. 1% of partial melt and is the main source of regional Quaternary volcanism [56].

Third, gas-water-rock-interactions, including dissolution of C- and Fe-bearing minerals in water at ~300 °C and carbonate methanation between 250 and 800 °C, do not depend on magma or magma-derived fluids [2,5]. The “Klerf Schichten” (Lower Ems) are alternating layers of reddish Fe-bearing sandstones and C-bearing shales and schists ≤ 2200-m thick and may be suitable formations for decomposition of C- and Fe-bearing minerals. Paleozoic bedrock sediments, especially the “Sphaerosiderith Schiefer” (Upper Ems; ≤ 150-m thick) schists with iron concretions (“Eisengallen”), are suitable formations for carbonate methanation: the decomposition of carbonate minerals (calcite, magnesite, siderite) at lower temperatures in H_2_-rich environments without mediation of gaseous CO_2_ (as it is usually the case for catalytic hydrogenation or FTT reaction) [2]. Within the habitable zone in the upper crust, at temperatures >150 °C and in the presence of CO_2_, CO, and H_2_, CH_4_ may be produced in aqueous solution even in the absence of a heterogeneous catalyst or gas phase by a series of redox reactions leading to the formation of formic acid, formaldehyde and methanol. Finally, abiotic CH_4_ also can form *in situ* through low temperature processes including the Sabatier and Fischer-Tropsch type (FTT) synthesis reactions with metals like Fe or Ni or clay minerals as catalysts [2,5].

Because the largest quantities of abiotic gases found on Earth’s surface are produced by low-temperature gas-water-rock reactions [54] we attribute the −37‰ δ^13^C-CH_4_ signature in RWA nests to fault-related emissions of abiotically formed CH_4_ by gas-water-rock reactions occurring at low-temperatures in a continental setting at shallow depths (micro-seepage). Probable sources might be Devonian schists (“Sphaerosiderith Schiefer”) with iron concretions (“Eisengallen”) sandstones and/or the iron-bearing “Klerf Schichten”. However, we cannot exclude the possibility of overlap by magmatic CH_4_ micro-seepage from the Eifel plume.

In summary, we suggest that RWA nests can indicate actively degassing faults trapping migrating CH_4_ from the deep underground, but that future work should seek to determine if the −37‰ signature can be attributed to a purely abiotic source, or a combination of abiotic/thermogenic source. Such a study should use additional measurements of δ^13^H and run long enough to determine the influence of irregularly timed earthquake events on patterns of methane degassing.

### Earth tides and earthquakes

Earth tides were basically semi-diurnal. Methane activity (Fig 8A, B) showed a low negative correlation with earth tides of ≈-0.4 at a lag of 6-8 hours. The cross-correlation between the earth tides and δ^13^C-CH_4_ was ≤ |0.15| (Fig 8C). Only one earthquake occurred nearby (local magnitude: 0.8; depth: km; distance: 20 km; Fig 4). This micro-earthquake neither influenced degassing nor RWA activity.

**Fig 8.**
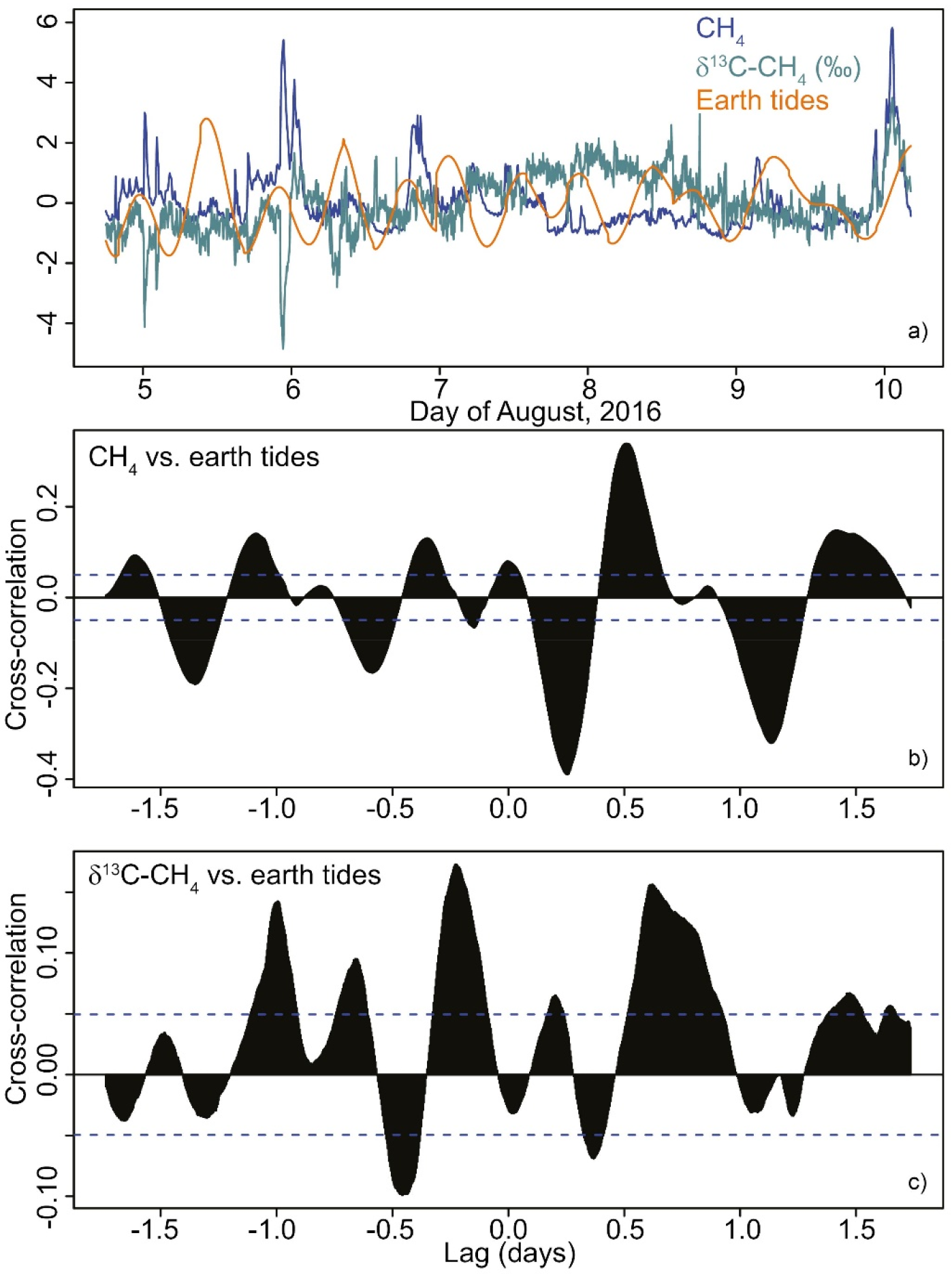
Relationships between CH_4_ (blue), δ^13^C-CH_4_ (green), and earth tides (orange). Time-series (a) of centered and scaled data. Cross-correlation of the time-series of CH_4_ (b) and δ^13^C-CH_4_ (c) with earth tides.

### RWA activities and external parameters

Neither our second or third hypotheses were supported by the data. During the investigation period, ant activity was higher than we had observed in 2009-2012, although an “M-shaped” pattern in daily activity was still identifiable [36]. Relatively high RWA activities during the late afternoon and early evening hours could be attributable to direct sun hitting the nest during that time or with activities associated with rebuilding damage to the nest that had occurred on 18 March. We did not find any evidence that ant activity changed during the CH_4_ (micro)-seepage process, or that there were strong effects of weather (see also 36]), or methane seepage. Additional external agents, including mice and “anting” birds, or micro-earthquakes did not influence ant activities during the sampling week. We conclude that during our 8-day sampling period, RWA activity was independent from external agents.

### Nest gas CH_4_ and δ^13^C-CH_4_ and external parameters

We also did not find strong support for a relationship between CH_4_ in the nest and external variables during our 8-day sampling period. Atmospheric CH_4_ concentrations were always lower than CH_4_ in the RWA nest and there seemed to be little influence of atmospheric CH_4_ on CH_4_ in the nest. Less than 25% of the variance in CH_4_ and δ^13^C-CH_4_ were accounted for by weather conditions (*cf*. [57]). Earth tides also were not correlated with methane degassing in the nest. The −37‰ δ^13^C-CH_4_ signature in nest gas was detected only once. The micro-earthquake on August 9 did not influence CH_4_ degassing because of its far distance (20 km). On August 13, there was another earthquake (ML: 0.7; D = 13 km) only 2.3 km away from the nest. It might be, that the −37‰ δ^13^C-CH_4_ signature in nest gas was a precursor to the August 13 earthquake, promoting degassing due to an increase in compressive stress [9,10]. But this remains unanswered as the CH_4_ measurement campaign was terminated at August 11.

## Conclusions

For the first time, both CH_4_ and δ^13^C-CH_4_ in a RWA nest was continuously recorded *in situ*. Methane degassing nor RWA activity was synchronized with earth tides, micro-earthquakes, or weather conditions. Elevated CH_4_ concentrations in nest gas appear to result from a combination of microbial activity and fault-related emissions moving via through fault networks through the RWA nest. Two δ^13^C-CH_4_ signatures were identified in nest gas: −69‰ and −37‰. The −69‰ signature of δ^13^C-CH_4_ within the RWA nest is best attributed to microbial decomposition of organic matter. This finding supports previous findings that RWA nests are hot-spots of microbial CH_4_. Additionally, the −37‰ δ^13^C-CH_4_ signature is the first evidence that RWA nests also may serve as traps for fault-related emissions of CH_4_. The −37‰ δ^13^C-CH_4_ signature can be attributed either to thermogenic/fault-related or to abiotic/fault-related CH_4_ formation originating from, e.g., low-temperature gas-water-rock reactions in a continental setting at shallow depths (micro-seepage). Future work on the −37‰ signature should use additional measurements of δ^13^H and run long enough to determine the influence of irregularly timed earthquake events on patterns of methane degassing.

## Acknowledgements

We thank Daniela Polag and Jan Hartmann (University of Heidelberg) for doing the nest-gas sampling. RWA activity recording was done using equipment from the Department of Geology at University of Duisburg-Essen. We also thank Dr. Peter Henrich (Leiter der Direktion Landesarchäologie - Außenstelle Koblenz) for his permission to conduct the survey on the Goloring site, and Hans-Toni Dickers, Paul Görgen and Bernd Klug from Kuratorium für Heimatforschung und -pflege, Kobern-Gondorf für their support during the field campaign.

## References

01. Keppler F, Boros M, Frankenberg C, Lelieveld J, McLeod A, Pirttilä AM, et al. Methane formation in aerobic environments. Env Chem. 2009;6(6):459–465. doi.org/10.1071/EN09137

02. Etiope G, Sherwood Lollar B. Abiotic methane on earth. Reviews of Geophysics. 2013;51. doi:10.1002/rog.20011.

03. Saunois M. et al. The Global Methane Budget: 2000-2012. Earth Syst. Sci. Data Discuss. 2016. doi:10.5194/essd-2016-25, 2016

04. Etiope G, Schoell M. Abiotic gas: atypical, but not rare. Elements – An International Magazine of Mineralogy, Geochemistry, and Petrology. 2014;10: 291–296. doi:10.2113/gselements.10.4.291.

05. Kiätävienen R, Purkamo L. The origin, source, and cycling of methane in deep crystalline rock biosphere. Front. Microbiol. 2015;6:725. doi:10.3389/fmicb.2015.00725.

06. Jílková V, Picek T, Šestauberová M, Krištůfek V, Cajthaml T, Frouz J. Methane and carbon dioxide flux in the profile of wood ant (Formica aquilonia) nests and the surrounding forest floor during a laboratory incubation. FEMS Microbiol Ecol. 2016;92:fiw141 doi.org/10.1093/femsec/fiw141.

07. Schoell M. The hydrogen and carbon isotopic composition of methane from natural gases of various origins. Geochimica et Cosmochimica Acta. 1980;44: 649–661. doi:10.1016/0016-7037(80)90155-6.

08. Keppler F, Hamilton JTG, Bra M, Röckmann T. Methane emissions from terrestrial plants under aerobic conditions. Nature. 2006;439: 187–191. doi:10.1038/nature04420.

09. Birdsell DT, Rajaram H, Dempsey D, Viswanathan HS. Hydraulic fracturing fluid migration in the subsurface: a review and expanded modeling results. Water Resour. Res. 2015;51: 7159–7188.

10. Boothroyd IM, Almond S, Worrall F, Davies RJ. Assessing the fugitive emission of CH_4_ via migration along fault zones – Comparing potential shale gas basins to non-shale basins in the UK. STOTEN. 2017;580: 412–424. doi.org/10.1016/j.scitotenv.2016.09.052.

11. Ciotoli G, Lombardi S, Zarlenga F. Natural leakage of helium from Italian sedimentary basins of the Adriatic structural margin. perspectives for geological sequestration of carbon dioxide. Advances in the Geological Storage of Carbon Dioxide. 2006; 191–202.

12. Etiope G. Natural emissions of methane from geological seepage in Europe. Atmos Environ. 2009;43:1430–1443.

13. Berberich G. Identifikation junger gasführender Störungszonen in der West-und Hocheifel mit Hilfe von Bioindikatoren. Doctoral dissertation. University of Duisburg-Essen. 2010; Available from: http://duepublico.uni-duisburg-essen.de/servlets/DerivateServlet/Derivate-25819/Diss_Berberich.pdf.

14. Berberich G, Schreiber U (2013) GeoBioScience: Red Wood Ants as Bioindicators for Active Tectonic Fault Systems in the West Eifel (Germany). Animals. 2013;3: 475–498. doi:10.3390/ani3020475.

15. Berberich G, Grumpe A, Berberich M, Klimetzek D, Wöhler C. Are red wood ants (Formica rufa-group) tectonic indicators? A statistical approach. Ecol Ind. 2016;61: 968–979. doi:10.1016/j.ecolind.2015.10.055.

16. Del Toro I, Berberich GM, Ribbons RR, Berberich MB, Sanders NJ, Ellison AM. Nests of red wood ants (Formica rufa-group) are positively associated with tectonic faults: a double-blind test. Peer J. 2017. https://doi.org/10.7717/peerj.3903

17. Risch AC, Jurgensen MF, Schütz M, Page-Dumroese DS. The contribution of red wood ants to soil C and N pools and CO_2_ emissions in subalpine forests. Ecology. 2005;86: 419e430.

18. Risch AC, Schütz M, Jurgensen MF, Domisch T, Ohashi M, Finér L. CO_2_ emissions from red wood ant (Formica rufa group) mounds: Seasonal and diurnal patterns related to air temperature Ann. Zool. Fennici. 2005;42: 283–290.

19. Ohashi M, Finér L, Domisch T, Risch AC, Jurgensen MF. CO_2_ efflux from a red wood ant mound in a boreal forest. Agricultural and Forest Meteorology. 2005; 30: 131e136.

20. Ohashi M, Finér L, Domisch T, Risch AC, Jurgensen MF, Niemelä P. Seasonal and diurnal CO_2_ efflux from red wood ant (Formica aquilonia) mounds in boreal coniferous forests. Soil Biology & Biochemistry. 2007;39: 1504e1511.

21. Wu H, Lu X, Wu D, Song L, Yan X, Liu J. Ant mounds alter spatial and temporal patterns of CO_2_, CH_4_ and N_2_O emissions from a marsh soil. Soil Biology & Biochemistry. 2013;57: 884e891

22. Bender MR, Wood CW. Influence of red imported fire ants on greenhouse gas emissions from a piedmont plateau pasture. Communications in Soil Science and Plant Analysis. 2003;34:1873e1889.

23. Crockett RGM, Perrier F, Richon P Spectral-decomposition techniques for the identification of periodic and anomalous phenomena in radon time-series. Nat Hazards Earth Syst Sci. 2010;10: 559–564.

24. Etiope G, Klusman RW. Geologic emissions of methane to the atmosphere. Chemosphere. 2002;49:777–89.

25. Litt T, Brauer A, Goslar T, Merk J, Balaga K, Mueller H, et al. Correlation and synchronisation of Lateglacial continental sequences in northern Central Europe based on annually laminated lacustrine sediments. In: Bjorck, S, Lowe, J. J. and Walker M. J. C. editors. Integration of Ice Core, Marine and Terrestrial Records of Termination 1 from the North Atlantic Region. Quaternary Science Reviews. 2001;20:1233–1249.

26. LGB RLP. Geologie von Rheinland-Pfalz. Schweizbart’sche Verlagsbuchhandlung. Stuttgart. 2005.

27. Ritter JRR, Jordan M, Christensen U, Achauer U. A mantle plume below the Eifel volcanic fields, Germany. Earth Planet. Sci. Lett, 2001;186: 7–14.

28. Walker KT, Bokelmann GHR, Klemperer SL, Bock G Shear-wave splitting around the Eifel hotspot: Evidence for a mantle upwelling. Geophys J Int. 2005;163: 962–980.

29. Tesauro M, Hollenstein C, Egli R, Geiger A, Kahle HG. Analysis of central western Europe deformation using GPS and seismic data. J. Geodynam. 2006;42:194–209.

30. Clauser C, Griesshaber E, Neugebauer HJ. Decoupled thermal and mantle helium anomalies: Implications for the transport regime in continental rift zones. J Geophys Res. 2002;107: NO. B11, 2269, doi:10.1029/2001JB000675.

31. Hinzen KG. Stress field in the Northern Rhine area, Central Europe, from earthquake fault plane solutions. Tectonophysics. 2003;377: 325–356.

32. Dèzes P, Schmid SM, Ziegler PA. Evolution of the European Cenozoic Rift System: interaction of the Alpine and Pyrenean orogens with their foreland lithosphere. Tectonophysics. 2004;389: 1–133.

33. Ahorner L. Historical seismicity and present-day microearthquake activity in the Rhenish Massif, Central Europe. In: Fuchs K, von Gehlen K, Mälzer H, Murawski H and Semmel A., editors. Plateau Uplift. The Rhenish Shield - A Case History. Springer-Verlag, Berlin, 1983. pp 198–221.

34. Ziegler PA., Dèzes P Crustal Evolution of Western and Central Europe. Crust Europe 20.03.05, 31 p. 2005.

35. BNS. Earthquake Data Catalogue. Department of Earthquake Geology of Cologne University. http://www.seismo.uni-koeln.de/catalog/index.htm. Accessed 01 September 2016.

36. Berberich G, Berberich M, Grumpe A, Wöhler C, Schreiber U. First Results of 2.5 Year Monitoring of Red Wood Ants’ Behavioural Changes and Their Possible Correlation with Earthquake Events. Animals. 2013;3: 63–84. doi:10.3390/ani3010063.

37. Maes F, Collignon A, Vandermeulen D, Marchal G, Suetens P. Multimodality image registration by maximization of mutual information. IEEE transactions on medical imaging. 1997;16.2: 187–198.

38. Dehant et al. and Milbert D (2016) Version 15.02.2016 (http://geodesyworld.github.io/SOFTS/solid.htm).

39. LGB RLP. Erdbebenereignisse lokal. Aktuelle Erdbebenereignisse in Rheinland-Pfalz, Baden-Württemberg und in 1000 km Entfernung. Landeserdbebendienst Rheinland-Pfalz. http://www.lgb-rlp.de/fachthemen-des-amtes/landeserdbebendienst-rheinland-pfalz/. Accessed 01 September 2016.

40. Kristan M, Leonardis, A, Skočaj, D. Multivariate online kernel density estimation with Gaussian kernels. Pattern Recognition. 2011. doi:10.1016/j.patcog.2011.03.019

41. Reimann C, Filzmoser P, Garrett RG. Background and threshold: critical comparison of methods of determination. Sci. Total Environ. 2005;346: 1–16.

42. Phillips NG, Ackley R, Crosson ER, Down A, Hutyra LR, Brondfield M, Karr JD, Zhao KG, Jackson RB. Mapping urban pipeline leaks: methane leaks across Boston. Environ. Pollut. 2013;73: 1–4.

43. Hinkle DE, Wiersma W, Jurs SG. Applied Statistics for the Behavioral Sciences. 5. ed., [reprint], internat. student ed., Wadsworth Publishing, Belmont, CA. 2009.

44. Pataki DE, Ehleringer JR, Flanagan LB, Yakir D, Bowling DR, Still CJ, et al. The application and interpretation of Keeling plots in terrestrial carbon cycle research. Global biogeochemical cycles. 2003;17: 1022. doi:10.1029/2001GB001850, 2003.

45. Etiope G, Vadillo I, Whiticar MJ, Marques JM, Carreira PM, Tiago I, et al. Abiotic methane seepage in the Ronda peridotite massif, southern Spain. Applied Geochemistry. 2016;66: 101–113.

46. Whiticar MJ. Carbon and hydrogen isotope systematics of bacterial formation and oxidation of methane. Chem Geol. 1999;161: 291–314.

47. Thielemann T, Lucke A, Schleser GH, Littke R. Methane exchange between coalbearing basins and the atmosphere: the Ruhr Basin and the Lower Rhine Embayment, Germany. Org Geochem 2000;31:1387–1408.

48. Alsaab D, Elie M, Izart A, Sachsenhofer RF, Privalov VA, Suarez-Ruiz I, Martinez L, Panova EA. Distribution of thermogenic methane in Carboniferous coal seams of the Donets Basin (Ukraine): “Applications to exploitation of methane and forecast of mining hazards”. Int J Coal Geol. 2009;78: 27–37.

49. EnergieAgentur NRW. Mine Gas. An energy source in Northrhine-Westphalia. EnergieAgentur NRW 1/2009.

50. Mills CT, Amano Y, Slater GF, Dias RF, Iwatsuki T, Mandernack KW. Microbial carbon cycling in oligotrophic regional aquifers near the Tono Uranium Mine, Japan as inferred from d13C and D14C values of in situ phospholipid fatty acids and carbon sources. Geochim Cosmochim Acta. 2010;74: 3785–3805.

51. Stępniewska Z., Kuźniar A. Endophytic microorganisms – promising applications in bioremediation of greenhouse gases. Appl Microbiol Biotechnol. 2013;97: 9589–9596. doi:10.1007/s00253-013-5235-9.

52. Stępniewska Z, Goraj W, Kuźniar A. Transformation of methane in peatland environments. Lesne Prace Badawcze (Forest Research Papers). 2014;75: 101–110. doi:10.2478/frp-2014-0010.

53. Horita J, Berndt ME. Abiogenic methane formation and isotopic fractionation under hydrothermal conditions, Science. 1999;285: 1055–7. doi:10.1126/science.285.5430.1055.

54. Etiope G, Ionescu A. Low-temperature catalytic CO_2_ hydrogenation with geological quantities of ruthenium: a possible abiotic CH_4_ source in chromititerich serpentinized rocks. Geofluids. 2015;15: 438–452. doi:10.1111/gfl.12106.

55. Etiope G, Judas J, Whiticar MJ. Occurrence of abiotic methane in the eastern United Arab Emirates ophiolite aquifer. Arab J Geosci. 2015;8: 11345–11348. doi10.1007/s12517-015-1975-4.

56. Ritter JRR. The Seismic Signature of the Eifel Plume. In: Mantle Plumes – A Multidisciplinary Approach. Eds: Ritter, Springer-Verlag Berlin Heidelberg, 2007. doi:10.1007/978-3-540-68046-8_12.

57. Toutain JP., Baubron JC. Gas geochemistry and seismotectonics: A review. Tectonophysics. 1999;304: 1–27.

58. Jackson RB, Down A, Phillips NG, Ackley RC, Cook CW, Plata DL, et al. Natural gas pipeline leaks across Washington, DC. Environ. Sci. Technol. 2014;48: 2051–2058.

59. Zazzeri G, Lowry D, Fisher RE, France JL, Butler D, Lanoisellé M, et al. Identification of urban gas leaks and evaluation of methane emission inventories using mobile measurements. Geophysical Research Abstracts. 2017; Vol. 19, EGU2017–14409.

